# Robust phylogenetic position of the enigmatic hydrozoan *Margelopsis haeckelii* revealed within the family Corymorphidae

**DOI:** 10.1101/2021.03.30.437375

**Authors:** Daria Kupaeva, Tatiana Lebedeva, Zachariah Kobrinsky, Daniel Vanwalleghem, Andrey Prudkovsky, Stanislav Kremnyov

## Abstract

The life-cycle and polyp morphology of representatives of Margelopsidae are very different from all other species in the hydrozoan clade Aplanulata. Their evolutionary origin and phylogenetic position has been the subject of significant speculation. A recent molecular study based only on COI data placed Margelopsidae as a sister group to all Aplanulata, an unexpected result because margelopsid morphology suggests affiliation with Tubulariidae or Corymorphidae. Here we used multigene analyses, including nuclear (18S rRNA and 28S rRNA) and mitochondrial (16S rRNA and COI) markers of the hydroid stage of the margelopsid species *Margelopsis haeckelii* Hartlaub, 1897 and the medusa stage of *Margelopsis hartlaubii* Browne, 1903 to resolve their phylogenetic position with respect to other hydrozoans. Our data provide strong evidence that *M. haeckelii*, the type species of *Margelopsis*, is a member of the family Corymorphydae. In contrast, *M. hartlaubii* Browne, 1903 is sister to *Plotocnide borealis* Wagner, 1885, a member of Boreohydridae. These results invalidate the family Margelopsidae. The phylogenetic signal of polyp and medusa stages is discussed in light of concept of inconsistent evolution and molecular phylogenetic analysis.

## Introduction

Species in the family Margelopsidae Mayer, 1910 (Aplanulata, Hydrozoa, Cnidaria) have intriguing life histories. The family is exclusively represented by hydrozoans with holopelagic life-cycles, where medusae and solitary vasiform polyps float freely throughout the water column. Interestingly, siphonophore specialists used margelopsid species as a model to explain the origin of siphonophoran colonies (Totton and Bargmann, 1965). Margelopsidae is comprised of three genera; *Margelopsis* Hartlaub, 1897; *Pelagohydra* Dendy, 1902; and *Climacocodon* Uchida, 1924, none of which have been sampled for comprehensive molecular analyses. Phylogenetic analysis using only COI sequences (Ortman et al, 2010) of *Margelopsis hartlaubii* Browne, 1903 suggested that Margelopsidae might be the sister group to the rest of Aplanulata. However, authors have not recovered strong support for this placement (Nawrocki et al., 2013). The systematics and phylogenetic position of Margelopsidae is solely based on insufficient morphological data. Given their polyp morphology, species of Margelopsidae show affinities with Tubulariidae or Corymorphydae, but their unique medusa morphology was used to justify their original erection as a separate family. Thus, sampling with more DNA markers and specimens – especially including the type species *Margelopsis haeckelii* – has been needed to determine the scope and phylogenetic position of the family Margelopsidae.

Despite difficulties of sampling margelopsid hydroids, we were finally able to collect representatives of *Margelopsis haeckelii* Hartlaub, 1897 and *Margelopsis hartlaubii* Browne, 1903 for molecular studies. *Margelopsis haeckelii* is the most studied species of its family, yet, documented collection records and morphological examinations have been very few (Hartlaub, 1897; Hartlaub, 1899; Lelloup, 1929; Werner; 1955, Schuchert, 2006). Polyps of *M. haeckelii* closely resemble tubulariid hydranths, having two whorls of tentacles but lacking both a hydrocaulus and stolonal system (Fig. 1, A, B). Free-swimming medusae develop from medusa buds located between whorls of polyp tentacles (Fig. 1, B, C, D). Eggs of *M. haeckelii* develop on the manubrium of the medusa (Fig. 1, C, D) and transform directly or through an encysted stage into a hydranth that never fixes to a substrate, exhibiting a continuous planktonic lifestyle (Werner; 1955). It is thought that eggs of this species are parthenogenetic, as no male gonads have ever been reliably documented. There is less information about *M. hartlaubii*, which is only known from the medusa stage. The medusa of *M. hartlaubii* can readily be distinguished from the medusa of *M. haeckelii* by its thick apical mesoglea of the bell without apical canal and two tentacles per bulb (Fig. 1,C, D, E) (Schuchert, 2006).

**Fig. 1.**
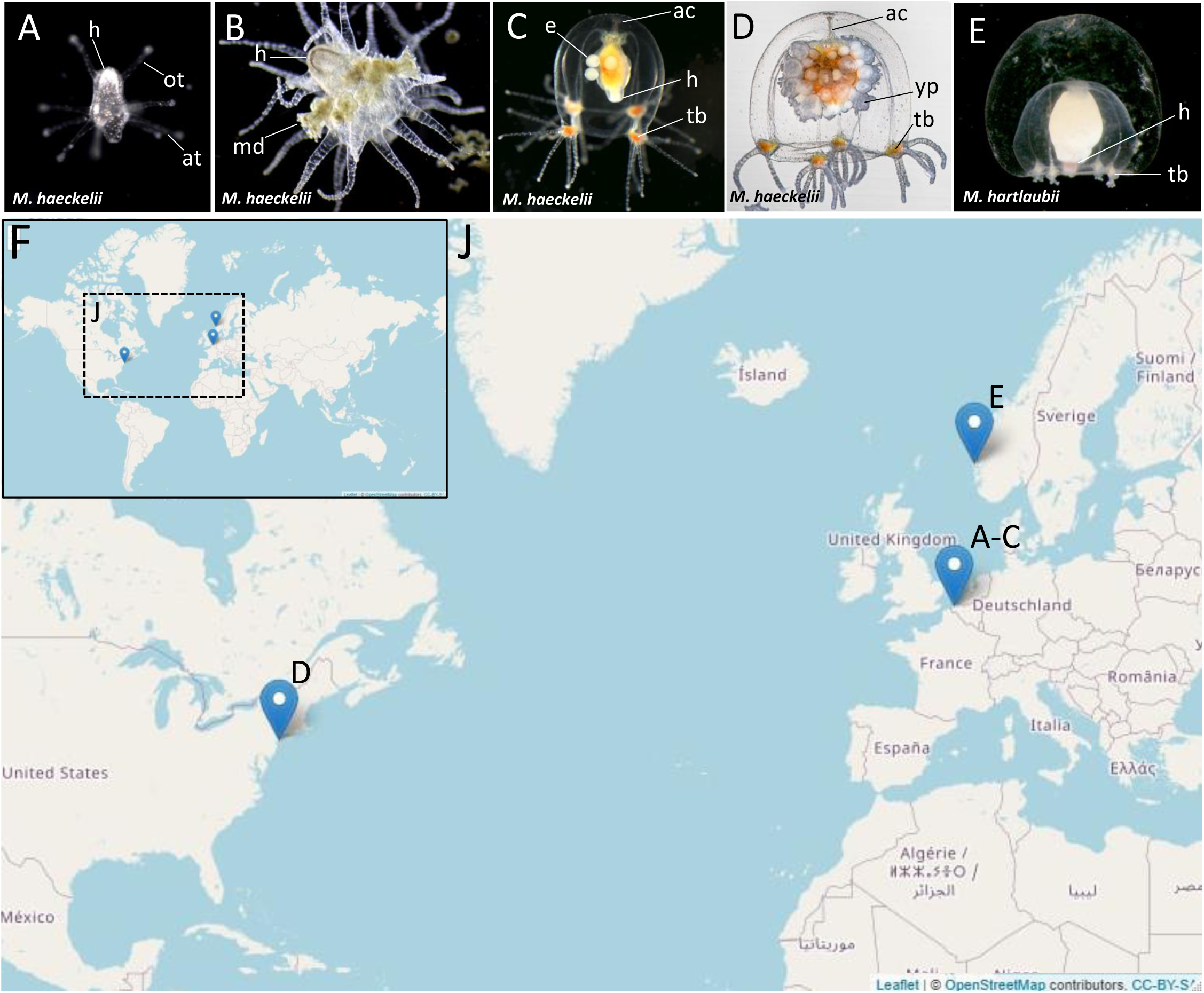
Morphology of collected Margelopsidae representatives and the locations of its samplings. (A-D) *Margelopsis haeckelii* Hartlaub, 1897. (A) Newly hatched polyp, (B) Mature polyp with medusa buds, (C, D) Mature medusa. (E) Mature medusa of *Margelopsis hartlaubii* Browne, 1903. Photo Credit: Dr. Peter Schuchert (Schuchert, 2022). (F, J) Geographic locations of sampling sites. Abbreviations, ac – apical canal, at – aboral tentacles, e – embryos, h – hypostome, md – medusoid ot – oral tentacles, tb – tentacle bulb, yp – young polyp.

In our study we obtained full-length sequences of 18S rRNA and 28S rRNA and partial sequences of the mitochondrial ribosomal 16S rRNA and cytochrome oxidase subunit I (COI) in order to phylogenetically place *M. haeckelii* and *M. hartlaubii* within as comprehensive sampling of hydrozoan taxa as possible. Using this approach, we provide the first molecular evidence that *M. haeckelii* should be placed within the family Corymorphydae. Our findings further showed that the previously sequenced *M. hartlaubii* is a relative of the family Boreohydridae, and is only distantly related to *Margelopsis haeckelii*, the type species of the genus.

## Methods and materials

### Animal sampling

Some *M. haeckelii* polyps were collected in the North Sea (loc. Belgium, Ostend, 51.218028°, 2.879417°) (Fig. 1,F, J). Polyps were collected with a plankton net in the coastal area. Collected animals were used to set up a lab culture. The obtained culture was maintained throughout the year in aquaria using artificial sea water (salt Red Sea Coral Pro, salinity 30–32‰) at the Department of Embryology, Lomonosov Moscow State University, Russia, Moscow. For both polyp and medusa stages, *Artemia salina* nauplii, at least 3 days after hatching, were used for feeding. Animals were fed once a day.

Also, *M. haeckelii* medusae were collected in the Atlantic Ocean, Atlantic Coast of North America (loc. USA, New York, 40.560556°, -73.882333°). Medusae were collected with a plankton net in the coastal area, about 10 meters out from the shore. Collected animals were fixed and stored in 96% ethanol (Fig. 1,F, J).

*M. hartlaubii* DNA was a gift from Dr. Peter Schuchert (Schuchert, 2022). The medusa was collected in Norway, Raunefiord (60.2575°, 05.1393°) with a plankton net from 200 to 0 m depth on 14-JUN-2016.

Meiobenthic polyps of *Plotocnide borealis* (formerly known as *Boreohydra simplex*; Pyataeva et al., 2016) were collected in the White Sea near the N.A. Pertsov White Sea Biological Station of the Moscow State University, Kandalaksha Bay, Russia (66.528056°, 33.185556°). Fine mud with polyps was collected with a light hyperbenthic dredge from depth 20-40 m. Collected individuals were fixed and stored in 96% ethanol.

### Identification of COI, 16S rRNA, 18S rRNA and 28S rRNA sequences

COI, 16S rRNA, 18S rRNA and 28S rRNA sequence fragments were amplified from genomic DNA using PCR methods. Genomic DNA was extracted using standard phenol/chloroform protocols. This method involved tissue digestion with proteinase K (20 mg/mL) in a lysis buffer (20 mM Tris-CL pH 8.0, 5 mM EDTA pH 8.0, 400 mM NaCl, 2%SDS), extraction with phenol/chloroform (1:1), precipitation with 0.1 vol 3M Sodium acetate and 1 vol. 100% Isopropanol and elution in mQ water.

For amplification, we used the following primers pairs:

16SAR (TCGACTGTTTACCAAAAACATAGC) and 16SBR (ACGGAATGAACTCAAATCATGTAAG) for 16S rRNA (Cunningham and Buss, 1993); and jGLCO1490 (TITCIACIAAYCAYAARGAYATTGG) and jGHCO2198 (TAIACYTCIGGRTGICCRAARAAYCA) for COI (Geller et al., 2013). Amplification programs used for 16S rRNA and COI are as previously described in Prudkovsky et al., 2019.

18S-EukF (WAYCTGGTTGATCCTGCCAGT) and 18S-EukR (TGATCCTTCYGCAGGTTCACCTAC) for 18S rRNA (Medlin et al., 1988). F97 (CCYYAGTAACGGCGAGT), R2084 (AGAGCCAATCCTTTTCC), F1383 (GGACGGTGGCCATGGAAGT) and R3238 (SWACAGATGGTAGCTTCG) for 28S rRNA (Evans et al., 2008). Amplification programs used for 18S rRNA and 18S rRNA are as previously described in Evans et al., 2008.

Full-length 18S rRNA and 28S rRNA sequences of *M. haeckelii* from the North Sea were obtained from the reference transcriptome available in our laboratory. For transcriptome sequencing, total RNA was extracted from a mixture of various *Margelopsis* life and developmental stages. Total RNA extraction was conducted using the Zymo Research Quick-RNA MiniPrep Plus Kit according to the manufacturer’s instructions. Poly-A RNA enrichment, cDNA library construction and sequencing were carried out at Evrogen (Russia). The cDNA library was sequenced using the Illumina NovaSeq 6000 SP flow cell to produce with 150-bp paired-end reads. The high-quality reads were employed for the *M. haeckelii* transcriptome assembly with the SPAdes assembler (v.3.13.1) (Bankevich et al., 2012).

### Phylogenetic analyses

Nucleotide sequences were aligned using the MUSCLE algorithm in MUSCLE software (v3.8.31) (Edgar et al., 2004) and trimmed with the TrimAL tool (v.1.2rev59) (Capella-Gutiérrez et al., 2009). A heuristic approach “automated1” was used to select the best automatic method for trimming our alignments.

Phylogenetic analyses were performed using Maximum Likelihood methods in IQTree v.2.0-rc2 software (Minh, et al., 2020) according to the optimal models for each gene. Individual marker analyses and a concatenated gene analysis were performed. The best models of nucleotide substitution were chosen using ModelFinder (Kalyaanamoorthy et al., 2017). The GTR+F+I+G4 was found to be optimal for the COI dataset; GTR+F+I+G4 for 16S rRNA; TIM3+F+R3 for 18S rRNA; and TIM3+F+R5 for 28S rRNA. One thousand bootstrap replicates were generated for each individual analysis, as well as for the combined analysis.

The concatenated COI+16S+18S+28S alignment was constructed using Sequence Matrix (https://github.com/gaurav/taxondna). The concatenated dataset was analyzed using IQTree (v.2.0-rc2) with partitioned analysis for multi-gene alignments (Chernomor, et al., 2016). The set of selected species for concatenated analysis was chosen mainly according to Nawrocki et al. (2013) and considering the availability of individual gene sequences in GenBank for COI, 16S rRNA, 18S rRNA and 28S rRNA.

Trees were visualized in FigTree v1.4.4 and processed with Adobe Illustrator CC. No alterations were made to the tree topology or the branch lengths.

An approximately unbiased (AU) test (Hidetoshi, 2002) was performed using IQTree software for testing alternative phylogenetic hypotheses.

## Data availability

Sequences obtained in this study have been deposited in GenBank under the following accession numbers: *Margelopsis haeckelii* (OK129327, OK139084, OK142735, OK127861, ON391039, ON391070), *Margelopsis hartlaubii* (ON237369, ON237671, ON237710), *Plotocnide borealis* (OK110252).

## Results

Our phylogenetic investigation of phylogenetic affinities of species of Margelopsidae was conducted employing Maximum likelihood analysis for all single gene datasets as well as our final concatenated four-gene dataset (COI, 16S rRNA, 18S rRNA, 28S rRNA). All taxa used in our analysis are arranged taxonomically in Table 1. All *M. haeckelii* and *M. hartlaubii* sequences (COI, 16S rRNA, 18S rRNA, 28S rRNA) were newly generated for this study. *M. hartlaubii* had previously only had COI available on GenBank (Ortman et al., 2010). Maximum Likelihood bootstrapping (MLB) analysis of the concatenated dataset recovered a relatively well resolved tree and recovered Margelopsidae paraphyly. *M. hartlaubii* was recovered sister to *Plotocnide borealis* Wagner, 1885 (MLB=100), forming a clade that affiliate with the family Boreohydridae, a sister taxon to all other Aplanulata genera (MLB = 100) (Fig. 2). Each individual COI, 16S rRNA, 18S rRNA or 28S rRNA analysis also recovered a strong supported affiliation of *M. hartlaubii* within Boreohydridae (MLB = 100) (Fig. 2). At the same time, both *M. haeckelii* from different locations nested within the clade of the Corymorphidae (MLB=89). This clade comprised two subclades, each well supported, one for genus *Euphysa*, including the type species *Euphysa aurata* Forbes, 1848, and the other for *Corymorpha + M. haeckelii*, including the type species, *Corymorpha nutans* M. Sars (Fig 2). *M. haeckelii* is nested inside the clade *Corymorpha bigelowi* Maas, 1905, *Corymorpha nutans* M. Sars, 1835, *Corymorpha sarsii* Steenstrup, 1855 and *Corymorpha pendula* L. Agassiz (MLB=89). Clade *Euphysa+Corymorpha+M. haeckelii* was recovered to be the sister to Tubulariidae (MLB=85), which together with *Branchiocerianthus imperator* Allman, 1885 constitute the superfamily Tubularioidea. Tubularioidea is recovered as sister to Hydridae (MLB=91). General topology of our phylogenetic tree obtained in combined analysis coincides with the Aplanulata tree published by Nawrocki et al., 2013.

**Table 1.**
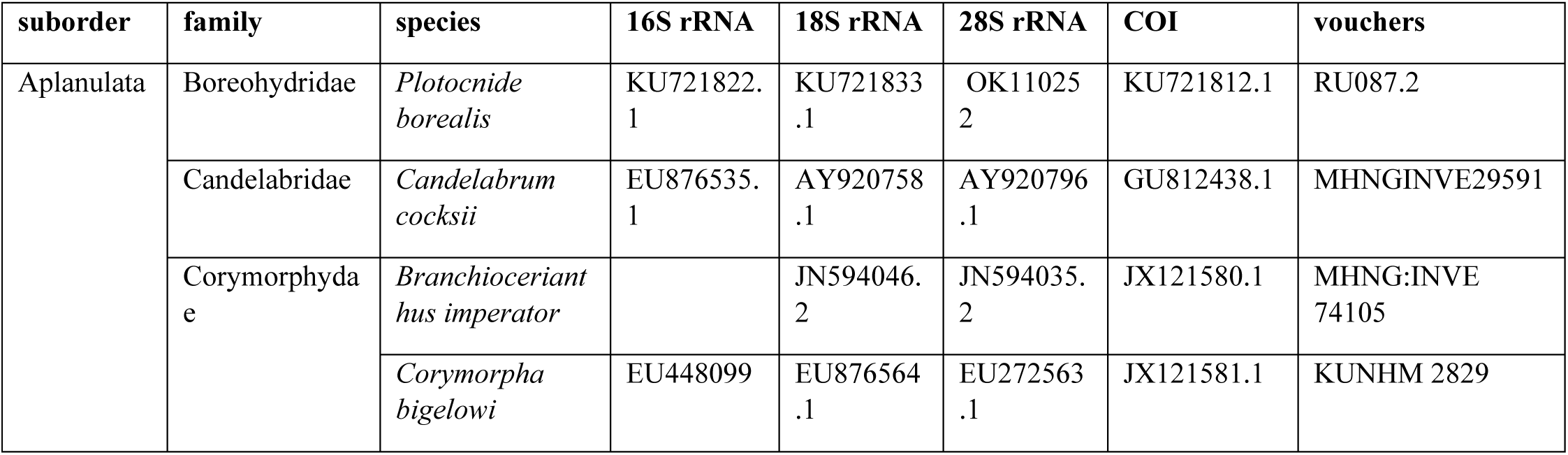
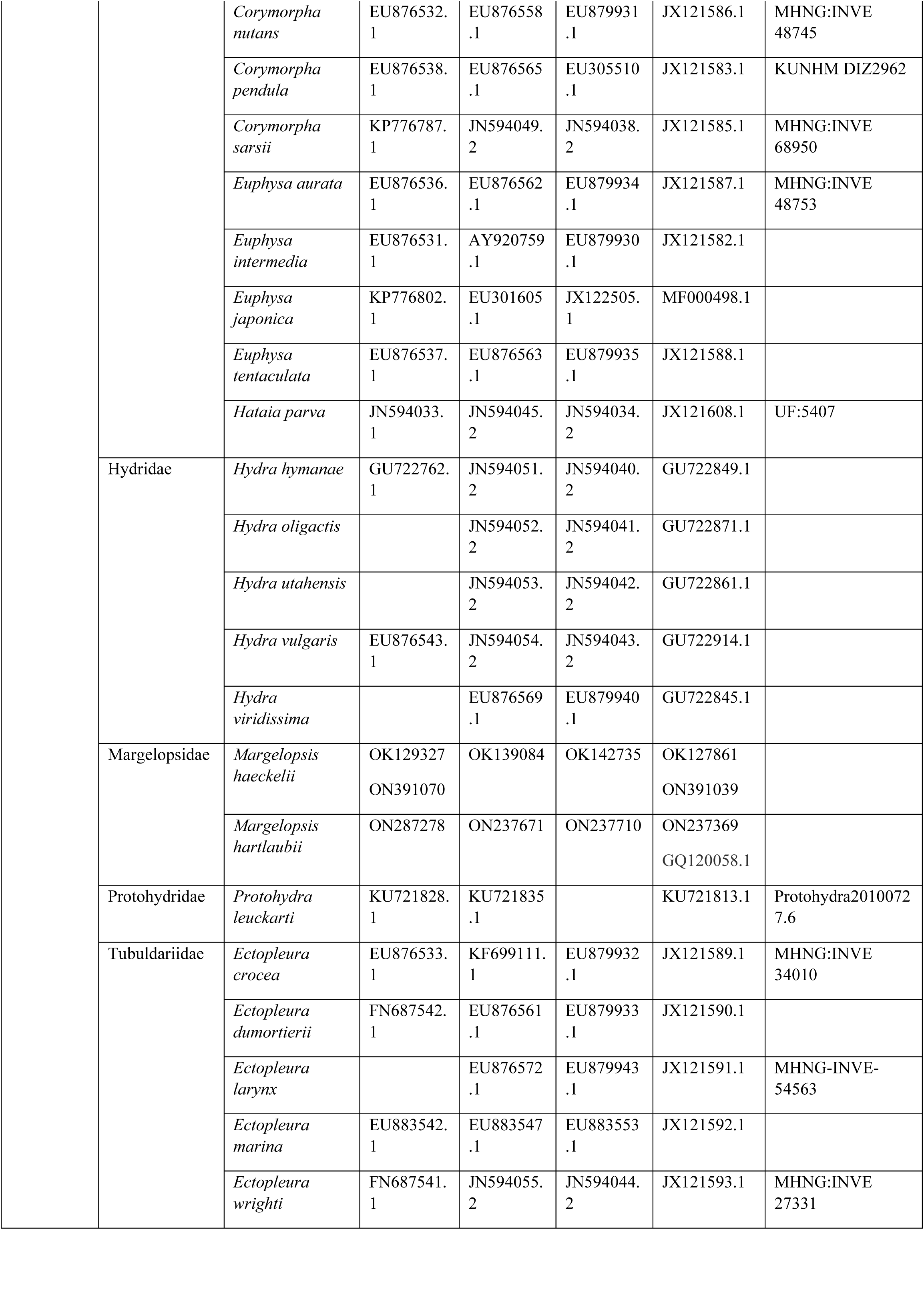
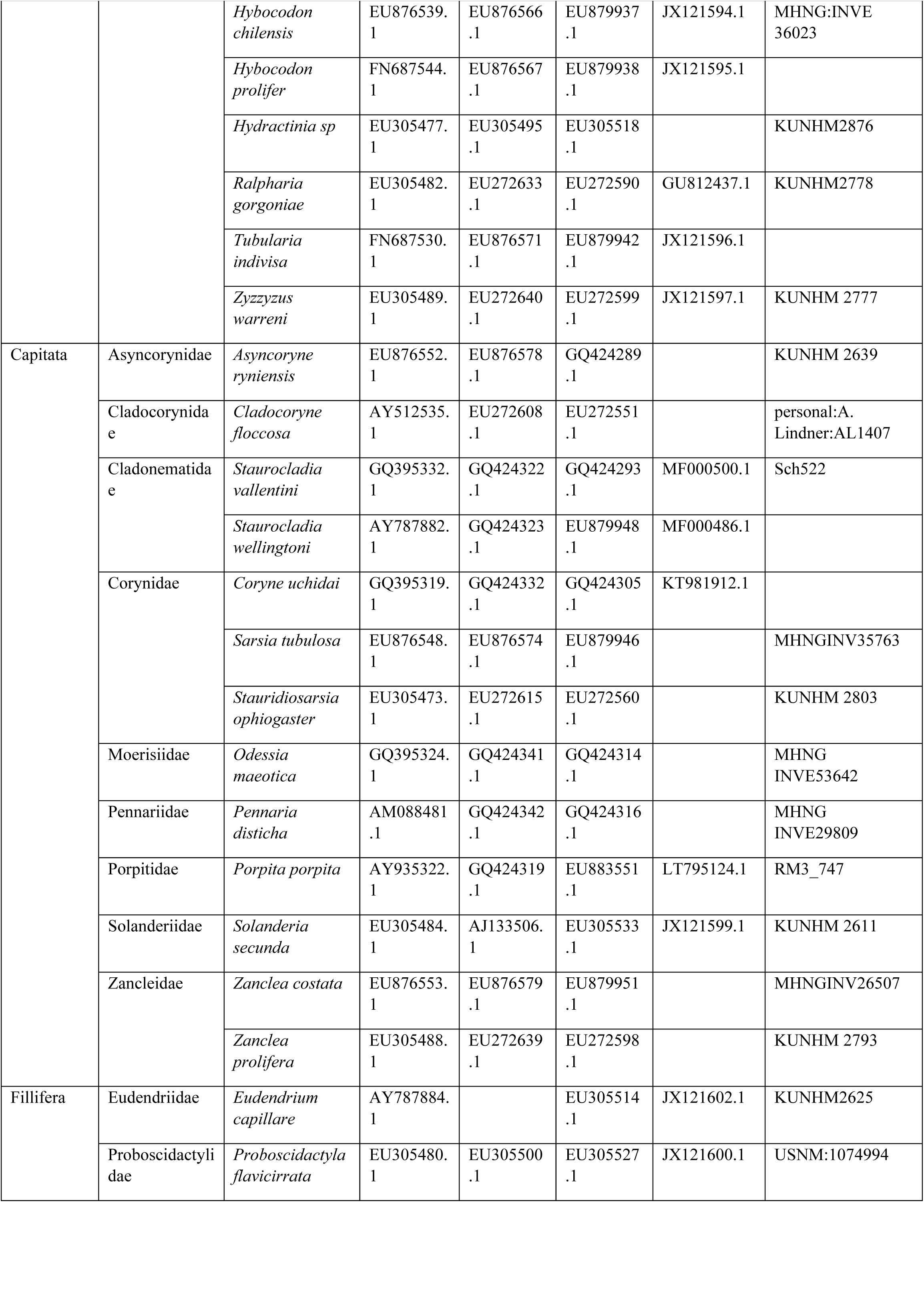
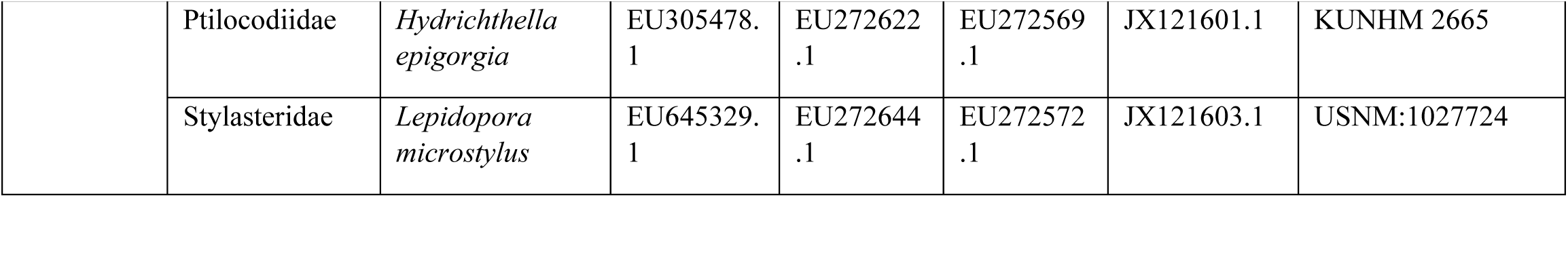
List of the species included in the study and corresponding GenBank accession numbers of all analyzed sequences.

**Fig. 2.**
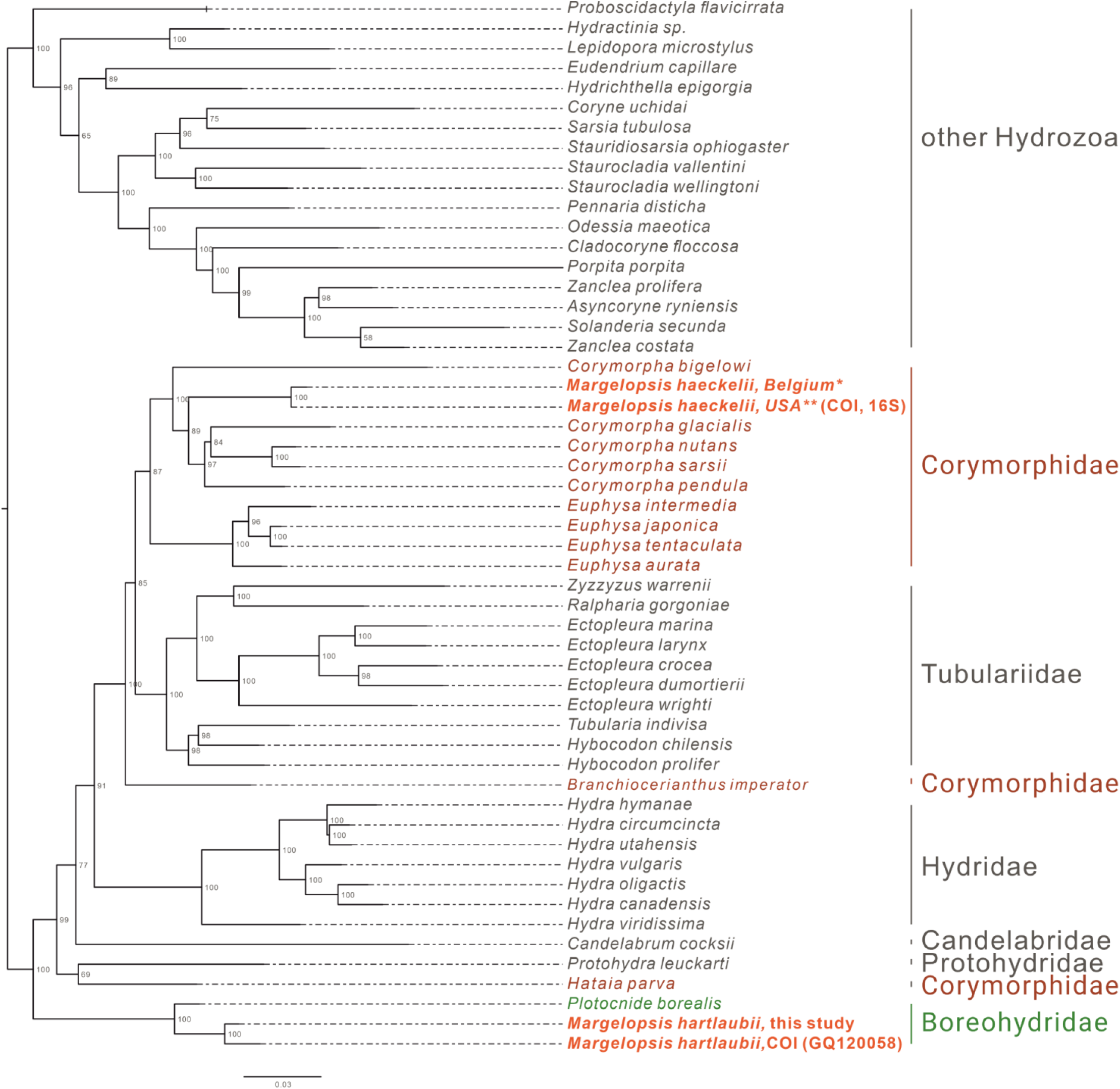
Analysis of phylogenetic position of *Margelopsis haeckelii* and *Margelopsis hartlaubii* in Aplanulata. Phylogenetic hypothesis of *Margelopsis haeckelii* relationships based on the combined mitochondrial and nuclear dataset (CO1+16S +18S+28S). Node values indicate bootstrap support from 1000 replicates. *Margelopsis haeckelii* and *Margelopsis hartlaubii* are in red. *WGS84 51.218028°, 2.879417°, ** WGS84 40.560556°, -73.882333°

Separate COI and 16S rRNA analysis recovered, that individuals of *Margelopsis haeckelii* from the opposite sides of the Atlantic Ocean are representatives of the same species (Fig. 1S, 2S). No nucleotide substitutions were identified in analyzed sequences of *Margelopsis haeckelii* from the waters of Belgium (51.218028°, 2.879417°) and the USA (40.560556°, -73.882333°).

At the same time, *M. hartlaubii* COI sequences analysis revealed five mismatches between sequences obtained in this study (ON237369) and sequence published in Ortman et al., 2010 (GQ120058.1) (Fig. 1S). However, COI sequences of *M. hartlaubii* published in Ortman et al., 2010 (GQ120058.1 and GQ120059.1) also are not identical and have three mismatches.

Phylogenetic hypothesis testing (AU test) was performed to test the statistical significance of tree topologies in our Maximum Likelihood analysis. The AU test rejected the phylogenetic hypothesis of the monophyly of *M. haeckelii* and *M. harlaubii*, providing strong evidence for the polyphyly of *Margelopsis*. Also, as our two individual marker analyses (16S and 28S) (Supp. 2, 3) placed *M. haeckelii* as a sister to *Corymorpha*, two hypotheses of alternative placements of *M. haeckelii* were evaluated: *M. haeckelii* is inside or outside *Corymorpha*. Results of the testing significantly support (p < 0.05) the hypothesis that *M. hackelii* is within *Corymorpha*. (Fig. 5S).

## Discussion

Our concatenated dataset (COI+16S+18S+28S), which included a comprehensive taxonomic sampling of hydrozoans, recovered *Margelopsis haeckelii* within Corymorphidae, nested within a clade consisting of several *Corymopha* species. This result is consistent with previous findings based solely on polyp morphology, where Margelopsidae was grouped with Tubulariidae and Corymorphidae in the superfamily Tubularoidea (Rees, 1957). Being quite small (1-2 mm), hydrocaulus-lacking pelagic polyps of the Margelopsidae are similar to those sessile polyps of corymophids and tubulariids despite the latter having a well-developed hydrocaulus and reaching up to ten centimeters in height. For all three families, hydranth tentacles are arranged into two, oral and aboral whorls and blastostyles are situated in the inter-tentacular region (Fig. 3, A, C). Our phylogenetic data support assertions that polyp tentacle patterns may be an important morphological character for identifying lineages in Aplanulata (Rees, 1957, Nawrocki et al. 2013).

**Fig. 3.**
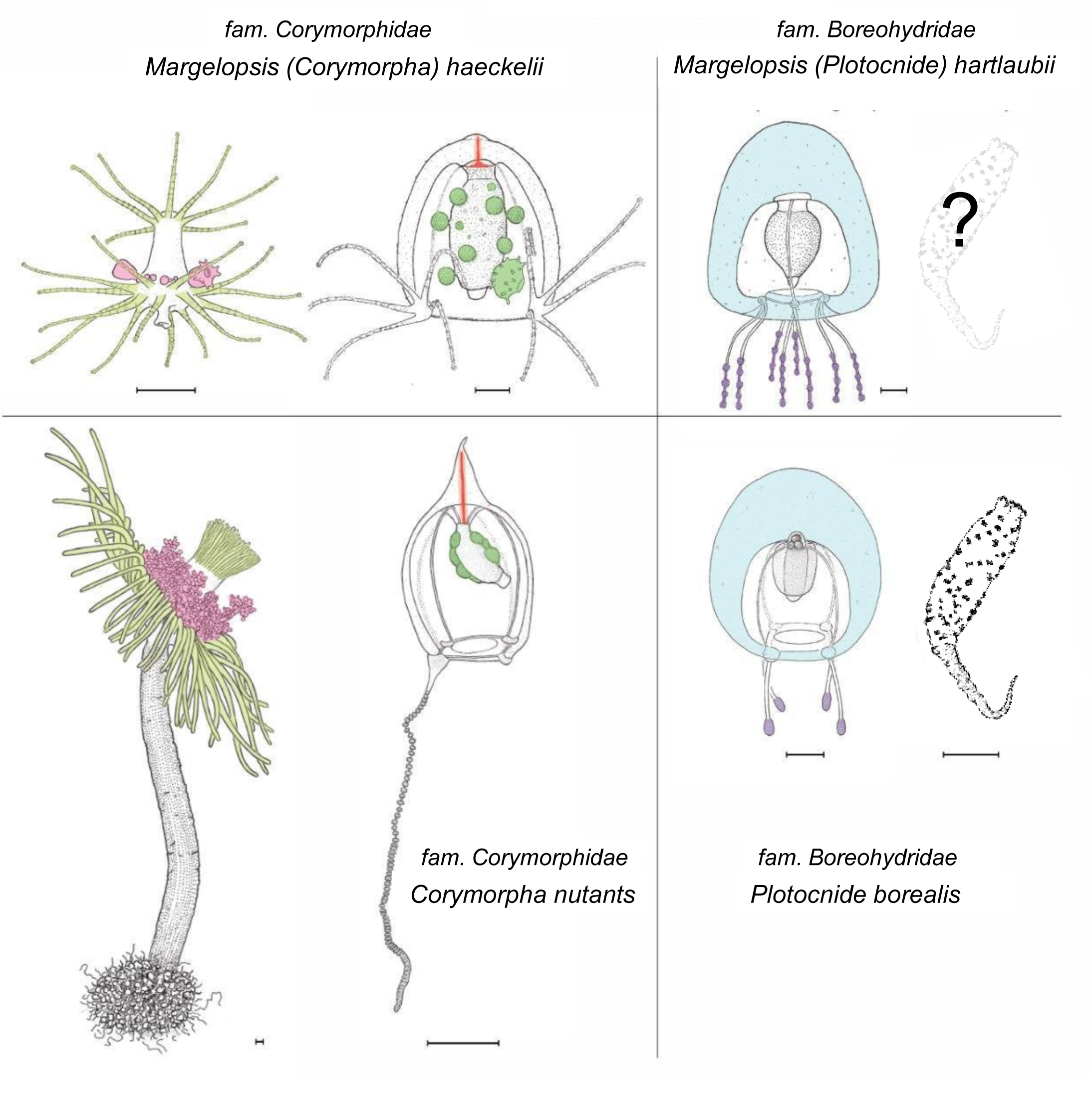
Comparison of morphological characters of (A) *Margelopsis hartlaubii*, (B) *Margelopsis haeckelii*, (C) *Corymorpha nutans* and (D) *Plotocnide borealis*. Scalebar – 0.4 mm. Color coding: yellow – oral and aboral whorls of polyp tentacles, pink– region of medusa budding, green – the region of gametes formation, orange – apical canal, blue – medusa umbrella with clusters of exumbrellar nematoblasts, violet – clusters of nematocysts located at the distal parts of tentacles. *Margelopsis hartlaubii, Margelopsis haeckelii, Corymorpha nutans* and *Plotocnide borealis* modified from Schuchert (2006; 2010)

Interestingly, *M. haeckelii* jellyfish are atypical in having radial symmetry, which more usually is bilateral in Aplanulata. The *M. haeckelii* jellyfish has 3-4 tentacles per bulb instead of one long tentacle per medusa, something typically seen among *Corymorpha* medusae. Even in *Euphysa*, the sister group to *Corymorpha*, radially symmetric adult medusae develop asymmetrically in contrast to medusae of *M. haeckelii*. The medusae of *Euphysa flammea* Hartlaub, 1902 only have a single tentacle in their youngest stage, with a second, third and fourth being added successively over time (Schuchert, 2010). Radially symmetric medusae in the species *P. borealis*, which is deeply nested in our phylogenetic analyses of Aplanulata (Pyataeva et al., 2016; this study) suggests that radial symmetry has re-evolved in *M. haeckelii*, a manifestation of the original body plan symmetry for medusae of Aplanulata. The presence of an apical canal in the umbrella may be a phylogenetically significant character warranting further investigation, as this character is shared both by *M. haeckelii* and all *Corymorpha* medusae (Fig. 3,A, C, marked orange). Reproductive characters appear to also reflect phylogenetic relationships in Aplanulata. Among all of Tubularoidea, only *Corymorpha* embryos undergo encystment similar to that of *M. haeckelii* (Petersen, 1990).

Surprisingly, our concatenated gene dataset, as well as our single gene COI dataset, recovered the medusa known as *M. hartlaubii* to be a close relative of *Plotocnide borealis*, and not closely related to *M. haeckelii* nor group within Corymorphidae. This result is further supported by independent morphological data showing several similarities between medusae of *M. hartlaubii* and *P. borealis*, including thick apical mesoglea of the bell (Fig. 3, marked blue), lack of an umbrella apical canal and nematocyst batteries being located at the distal parts of tentacles (Fig. 3, marked violet) (Schuchert, 2006). Based on our findings, medusae described by Browne (1903) have been wrongly attributed to the genus *Margelopsis*. Nawrocki et al. (2013) suggested that the hypothesis of *M. hartlaubii* as the sister to the rest of Aplanulata was uncertain due to low bootstrap support and that more genetic markers were needed to understand the phylogenetic placement of the species. Based on our multi-marker phylogenetic analysis and morphological data (Browne, 1903; Schuchert, 2006) we hypothesize that *M. hartlaubii* has a mud-dwelling, meiobenthic polyp like *P. borealis* (Fig. 3), and that the two species combined represent the sister group to the rest of Aplanulata.

In addition to *M. haeckelii* and *M. hartlaubii*, there are several other suspected species in the genus *Margelopsis*, including *Margelopsis gibbesii* (McCrady, 1859) and *Margelopsis australis* (Browne, 1910). Following Schuchert (2007), the World Register of Marine Species (https://www.marinespecies.org/) lists *Margelopsis gibbesii* as invalid. This stems from the fact that the original material used to describe this species, as *Nemopsis gibbesii*, consisted of a margelopsid polyp and a bougainvilliid medusa, the latter subsequently recognized as a medusa of *Nemopsis bachei* (L. Agassiz, 1862). This situation has generated subsequent nomenclatural confusion. More recently, Calder and Johnson (2015) stabilized the situation by designating the hydroid specimen illustrated by McCrady (1859) in Plate 10, Figure 7 as a lectotype for the margelopsid species. Calder and Johnson (2015) went on to provide evidence casting doubt on the distinction between M. gibbesii and M. haeckelii but maintained the two species given the geographic locations on either side of the north Atlantic and pending further study. In this study, however, using molecular phylogenetics, we have shown that *Margelopsis* from the western North Atlantic, and *M. haeckelii* from the eastern North Atlantic is the same species as *M. haeckelii, Margelopsis gibbesii* invalid. The lack of any nucleotide substitution in COI and 16S sequences of *Margelopsis* representatives from both sides of Atlantic Ocean makes it possible to suggest that these two populations are not isolated.

*Margelopsis australis* is only known from its original collection and is based on a single medusa specimen, lacking reliable characters for distinguishing it from *M. hartlaubii* (Browne 1910). Moreover, the single specimen was described as being “somewhat contracted and in a crumbled condition” (Browne 1910). Based on the available morphological data, we cannot state with any degree of certainty that *M. australis* is a valid species, or that it is a member of *Margelopsis*.

Medusae are a useful means of identifying species, genera and even family ranks (Rees, 1957; Bouillon, et al., 2006). A change in morphology of the typical jellyfish form within a family is usually due to the reduction of the medusa stage, something that is widespread throughout Anthoathecata and Leptothecata (Cornelius, 1992; Leclere et al., 2009; Cartwright, Nawrocki, 2010). However, *M. haeckelii* is a normally developed medusa, distinctly different from those typical of *Corymorpha*, despite their close relationship recovered by our phylogenetic analysis. Recent studies using molecular phylogenetic methods have revealed several such cases in which related taxa have very different jellyfishes or species with similar jellyfishes are only distantly related. The morphologically aberrant jellyfish *Obelia* is so different from other Companulariidae that a hypothesis was proposed for the re-expression of this jellyfish after its evolutionary reduction (Boero, Sara, 1987). However, this hypothesis was not supported by molecular phylogenetic analysis and *Obelia* may have originated from a *Clytia*-like ancestor (Cunha et al. 2017; Govindarajan et al., 2006; Leclere et al., 2019). *Larsonia pterophylla* (Haeckel, 1879) was previously assigned to the genus *Stomotoca* due to similarity of their jellyfishes (Larson, 1982). Interestingly, the structure of the polyps in the genera *Larsonia* and *Stomotoca* are so dissimilar that they could be attributed to different families (Boero, Bouillon, 1989). And indeed, according to molecular data, *L. pterophylla* and *Stomotoca atra* L. Agassiz, 1862 are not closely related. Rather, *L. pterophylla* is closely related to *Hydrichthys boycei* from the Pandeidae family, and *S. atra* is ungrouped with most species (Schuchert, 2018; Woodstock et al., 2019). Inclusion of the genus *Cytaeis* in Bougainvilliidae or the genera *Polyorchis* and *Scrippsia* in Corynidae is surprising due to the discrepancy between the jellyfishes of these genera and those typical of the respective families (Nawrocki, Cartwright, 2010; Prudkovsky et al., 2017). Finally, we conclude that appearance of atypical jellyfishes in hydrozoan families can indicate a great evolutionary plasticity of the medusa stage morphology. In contrast, the morphology of the hydroids appear to be more phylogenetically constant. For example, the morphology of *Cytaeis* hydroids is similar to the structure of Bougainvillidae hydroids with stolonal colonies, and Obelia-like polyps are typical for the family Campanulariidae (Prudkovsky et al., 2017; Leclere et al., 2019).

Concepts of ‘mosaic’ or ‘inconsistent evolution’ were proposed for these cases in which closely related hydroids can produce very different medusae or vice versa (Naumov 1956, 1960; Rees, 1957). Inconsistent evolution was explained by differences in the rate and direction of evolution in the two life cycle stages. Some incongruences between hydroid and medusa systems seem to result from weaknesses in a classification system (Petersen, 1990), but our work provides new reason to return to the discussion of this concept.

### Taxonomic recommendations

Based on our results, as well as a number of previous studies, we formally recommend the following changes to the taxonomy of Margelopsidae and its component species:

a. As multigene phylogenetic analyses nested *Margelopsis haeckelii*, the type species of *Margelopsis*, within genus *Corymorpha*, we recommend to redesignate it into *Corymorpha haeckelii*. *Corymorpha* M. Sars, 1835 Type species: *Corymorpha nutans* M. Sars, 1835 by monotypy. *Diagnosis*: Solitary hydroids with more or less vasiform hydranth, with long caulus or with short, squat polyp with broad head. **Rarely a hydrant without a caulus**. Hydranth with one or several closely set whorls of 16 or more **moniliform** or filiform tentacles and one or more aboral whorls of 16 or more long, non-contractile moniliform or filiform tentacles. Gastrodermal diaphragm parenchymatic or **without parenchymatic specializations of the gastrodermis**. Hydrocaulus, **if present**, stout, covered by thin perisarc, filled with parenchymatic gastrodermis, with long peripheral canals; aboral end of caulus with papillae turning more aborally into rooting filaments, rooting filaments scattered or gathered in a whorl, rooting filaments composed of epidermis and solid gastrodermis, sometimes tips with non-ciliated statocysts. **Otherwise, hydroid planktonic and hydrocaulus reduced, with a central depression**. With or without asexual reproduction through constriction of tissue from aboral end of hydrocaulus. Gonophores develop on blastostyles arranged in a whorl over aboral tentacles. Gonophores remain either fixed as sporosacs, medusoids, or are released as free medusae. Medusa bell apex dome-shaped or pointed, **with apical canal**. Four marginal bulbs present, lacking long exumbrellar spurs. With a single tentacle or three short tentacles and one long tentacle that differs not merely in size, but also in structure. **Rarely with 1-6 tentacle per bulb**. Manubrium thin-walled, sausage-shaped with flared mouth rim, reaching to umbrella margin. Cnidome comprises stenoteles, desmonemes, and haplonemes, **with or without euryteles**. *Remarks*: This diagnosis for the most part corresponds to Schuchert, 2010 (Schuchert, 2010), Petersen,1990 (Petersen, 1990) and Nawrocki et al., 2013 (Nawrocki et al., 2013), but with modifications (indicated in bold) to polyp and medusa body shape, and cnidome description to include *Margelopsis (Corymorpha) haeckelii*.
b. We suggest moving *Margelopsis hartlaubii* into family Boreohydridae and recommend to redesignate it into Plotocnide *hartlaubii*. *Plotocnide* Wagner, 1885 Type species: Plotocnide borealis Wagner, 1885 by monotypy. *Diagnosis:* Medusa umbrella evenly rounded with thick apical jelly and scattered groups of exumbrellar nematocysts; manubrium half as long as bell cavity, with or without broad, dome-shaped apical chamber; **without apical canal**; mouth simple, with ring of nematocysts; gonad forming thick ring around manubrium; four narrow radial canals and narrow ring canal; four marginal bulbs each **with 1-3 solid tentacles per bulb**; tentacles terminate in ovoid knob studded with nematocysts. No ocelli. Cnidome comprises desmonemes and stenoteles, **with or without mastigophores**. Hydroids, **if known**, solitary, small, with one whorl of reduced tentacles, capitate or not, located in the oral or median part of body; perisarc covering of base filmy or absent; gametes in body wall. Remarks: This diagnosis for the most part corresponds to Schuchert, 2006, 2010 (Schuchert, 2006; Schuchert, 2010), but with modifications (indicated in bold) to medusa body shape, and cnidome description to include *Margelopsis (Plotocnide) hartlaubii*.
c. We suggest that Margelopsidae should no longer be used, and both *Pelagohydra* and *Climacocodon* should be moved to Aplanulata *incertae sedis* until additional molecular phylogenetic analyses can clarify their phylogenetic placement.

## Conclusion

Our results clarify the phylogenetic picture of Aplanulata, by revealing the phylogenetic position of *M. haeckelii*, type species of the genus *Margelopsis* as falling within *Corymorpha* and *M. hartlaubii* as being a close relative of *Plotocnide* in the family Boreohydridae. On the case of the latter species, this phylogenetic result conflicts with the century old hypothesis that *Margelopsis* belongs to Tubulariidae or Corymorphidae (Nawrocki et al., 2013). However, by showing that *M. haeckelii* falls within the genus *Corymorpha*, our investigation presents strong evidence in support of this traditional hypothesis. Because *M. haeckelii* is a hydrozoan belonging to Corymorphidae, we can infer that this lineage evolutionarily lost their hydrocaulus and stolon, likely as an adaptation to a holopelagic life-cycle. It was previously suggested that the foundation for this type of changes in body plan, and accompanying life-style, might lead to speciation and could be reflected by changes in the expression of Wnt signaling components (Duffy, 2011). Based on our results, *M. haeckelii* might be a prime candidate for testing this hypothesis.

Unfortunately, due to the few and extremely irregular documented collection records of hydroids from the supposedly sister genera of *Margelopsis, Pelagohydra* and *Climacocodon*, the phylogenetic relationships within this group are still obscured. It remains unclear if Pelagohydra and Climacodon form a clade with either M. hartlaubii or M. haeckelii, or neither. Thus, the number of origin of a secondarily specialized pelagic polyp stage is still not known. The possible relationships between these three genera, as well as their phylogenetic placement, still need to be verified by additional studies when molecular data become available.

## Acknowledgements

We thank Dr. Peter Schuchert for the gift of *Margelopsis hartlaubii* DNA. We are grateful to Dr. Allen G. Collins for sequencing of the COI and 16S of *Margelopsis haeckelii* from NY (USA). We also thank Dr. Brett Gozales for the help with text and grammar editing. This study was supported by federal project 0088-2021-0009 of the Koltzov Institute of Developmental Biology of the Russian Academy of Sciences. This work was also supported by RSF, grant number 22-14-00116.

## Figure legends

**Fig. 1S.**
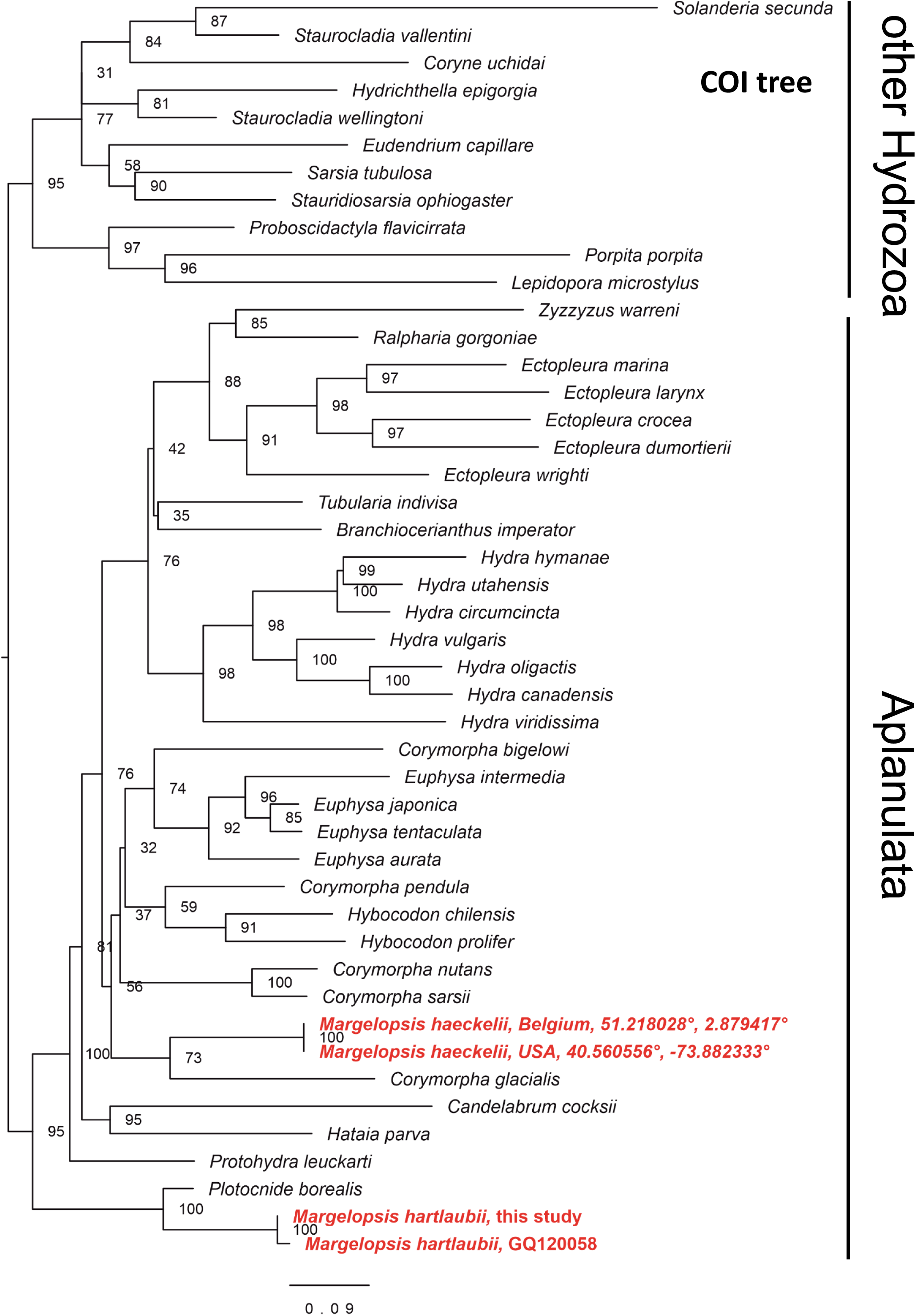
Phylogenetic hypothesis of *Margelopsis haeckelii* and *Margelopsis hartlaubii* relationships based on nuclear cytochrome oxidase subunit I (COI). Node values indicate bootstrap support from 1000 replicates. *Margelopsis haeckelii* and *Margelopsis hartlaubii* are in red.

**Fig. 2S.**
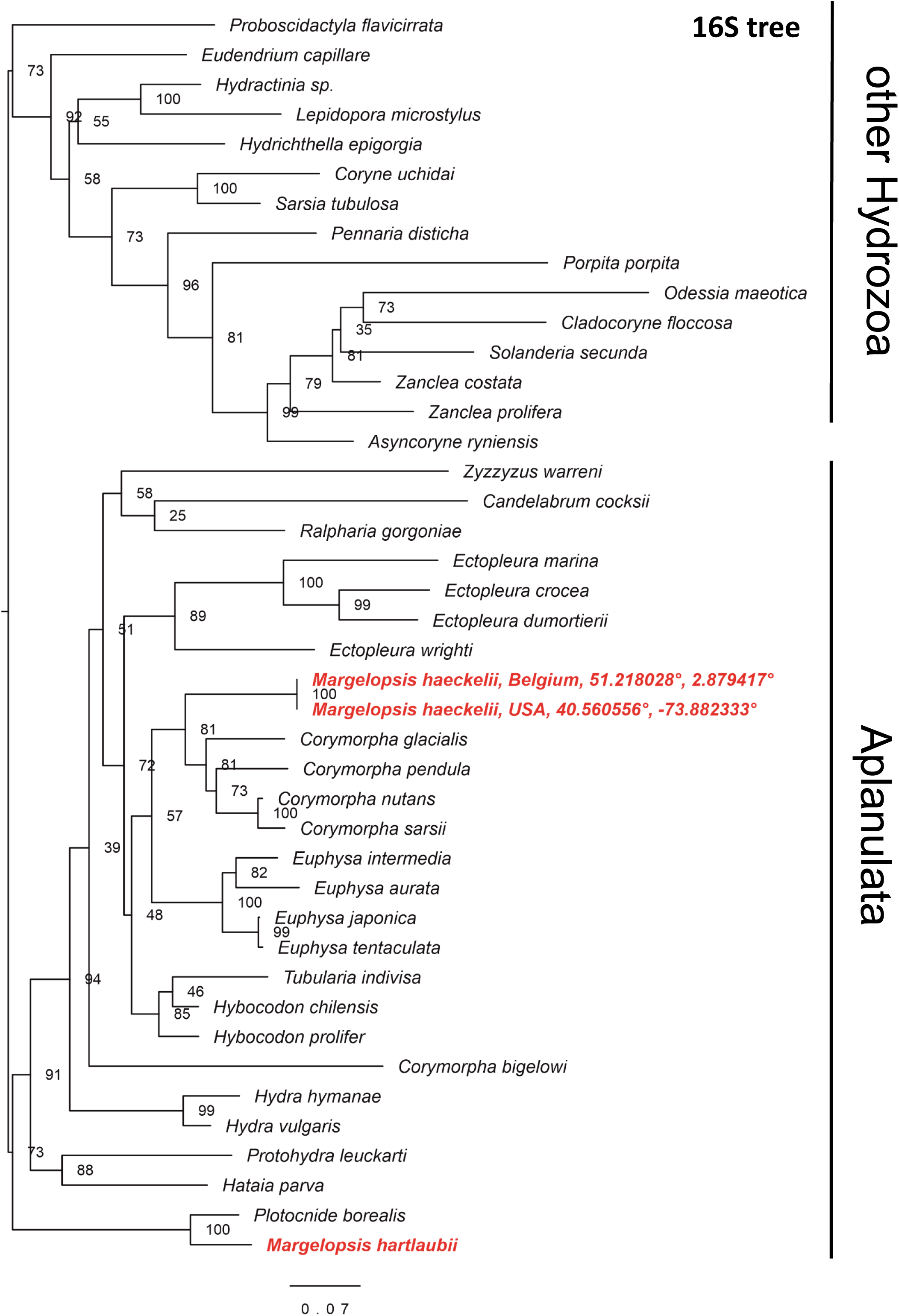
Phylogenetic hypothesis of *Margelopsis haeckelii* and *Margelopsis hartlaubii* relationships based on the mitochondrial 16S rRNA. Node values indicate bootstrap support from 1000 replicates. *Margelopsis haeckelii* and *Margelopsis hartlaubii* are in red.

**Fig. 3S.**
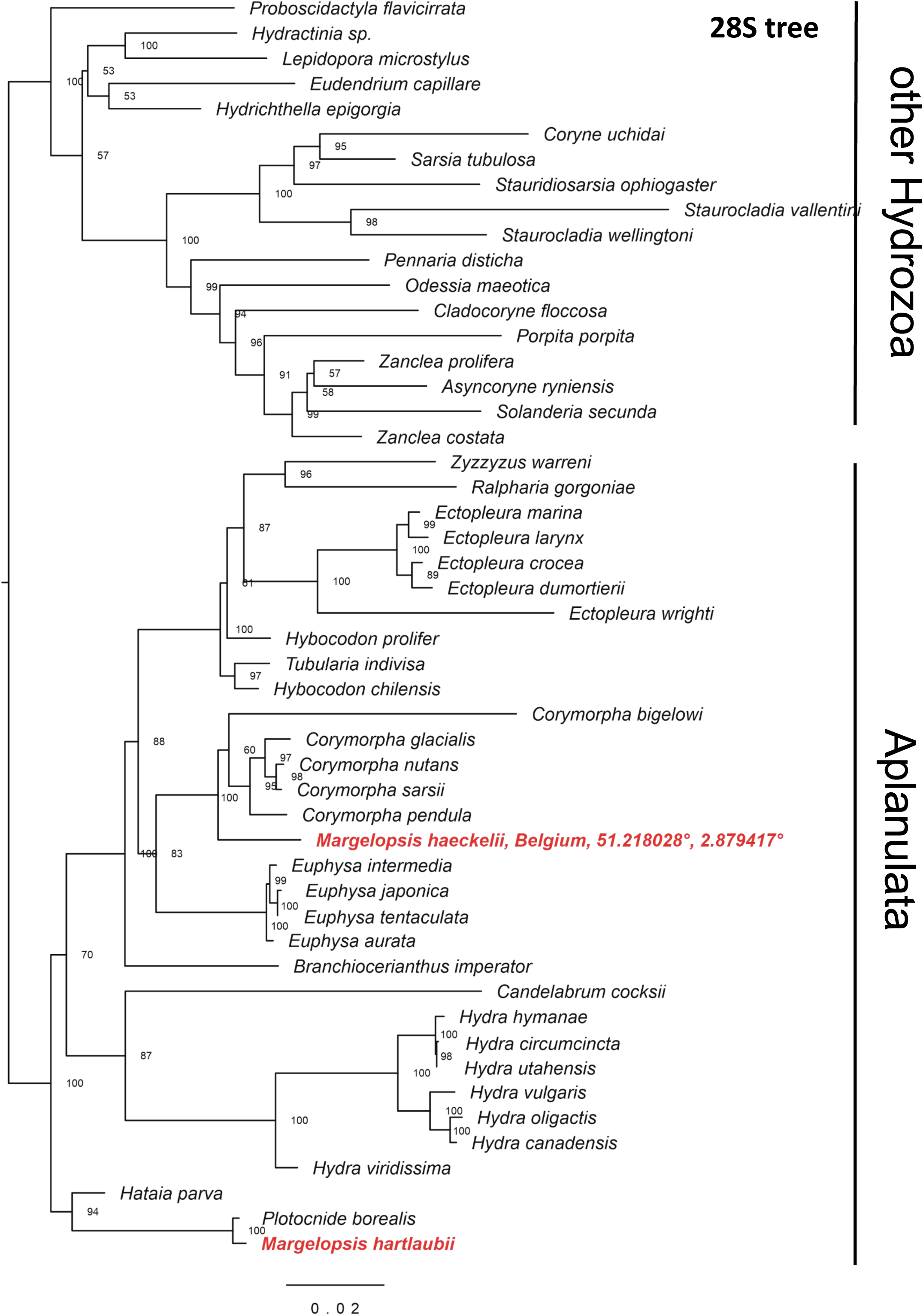
Phylogenetic hypothesis of *Margelopsis haeckelii* and *Margelopsis hartlaubii* relationships based on the 28S rRNA large ribosomal subunit. Node values indicate bootstrap support from 1000 replicates. *Margelopsis haeckelii* and *Margelopsis hartlaubii* are in red.

**Fig. 4S.**
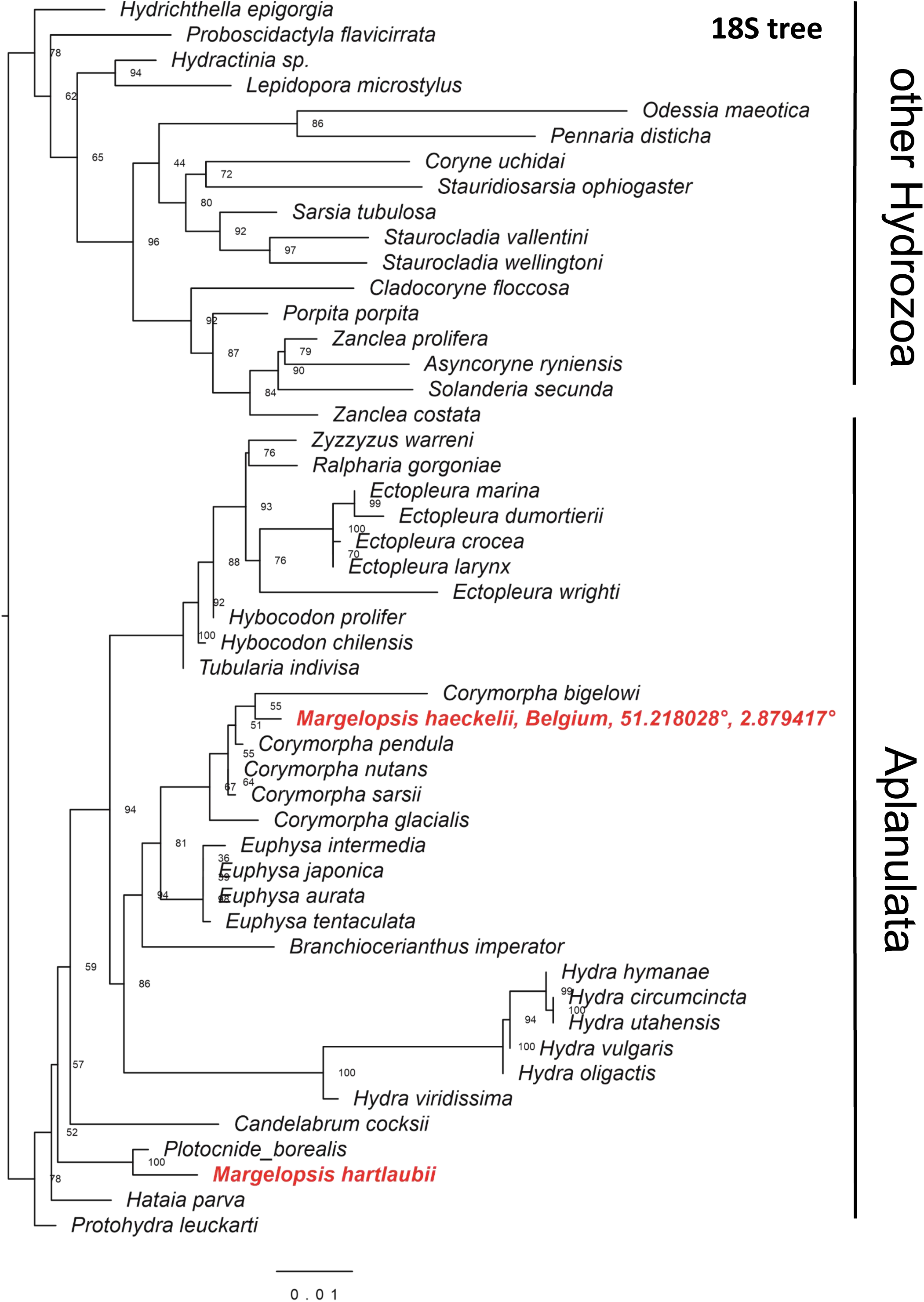
Phylogenetic hypothesis of *Margelopsis haeckelii* and *Margelopsis hartlaubii* relationships based on the 18S rRNA small ribosomal subunit. Node values indicate bootstrap support from 1000 replicates. *Margelopsis haeckelii* and *Margelopsis hartlaubii* are in red.

**Fig. 5S.**
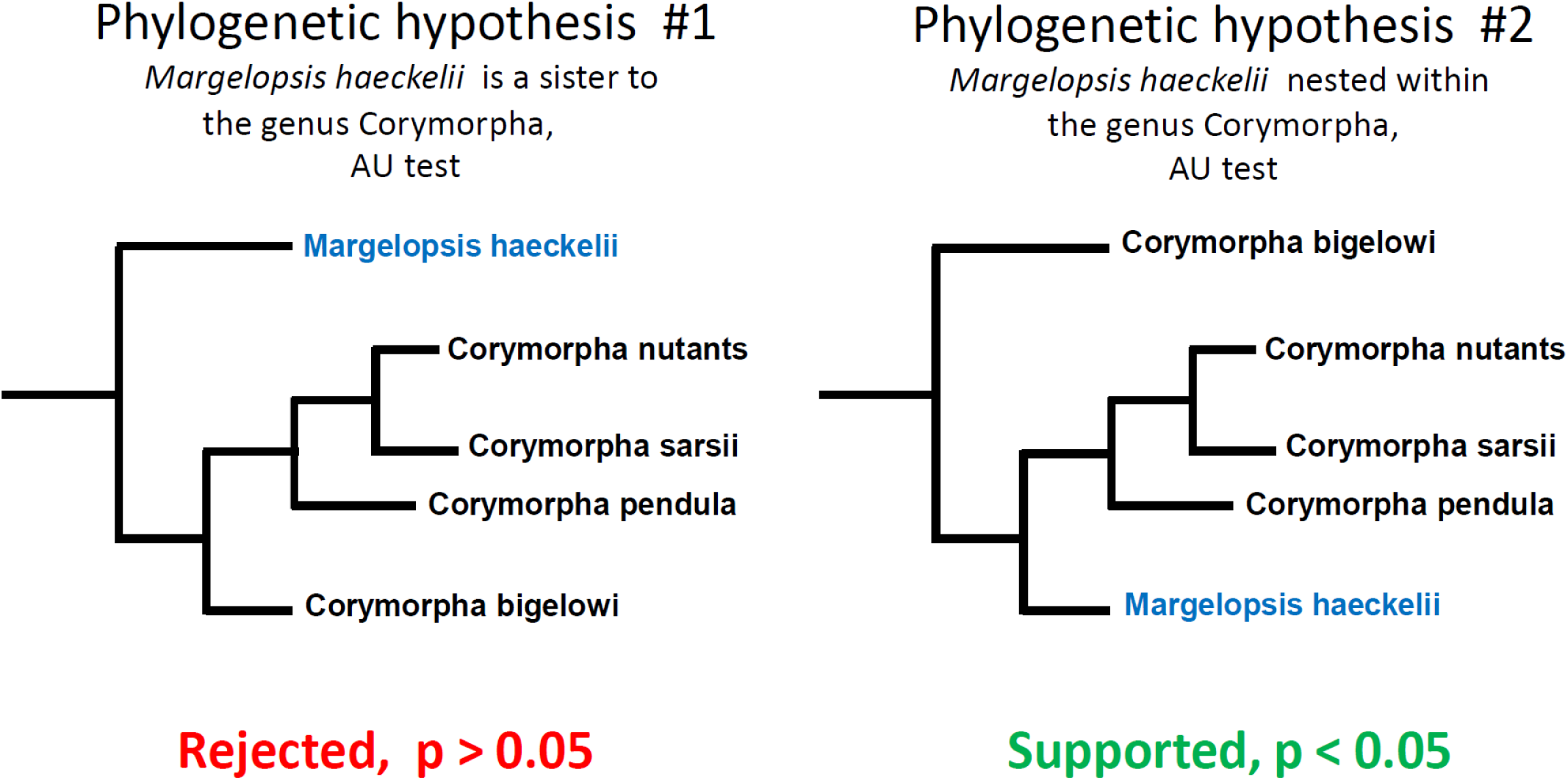
Testing of the phylogenetic hypotheses with AU test.

